# Understanding Fertility Intention Through the Lens of Family Functioning, Fertility Anxiety and Marriage-Fertility Views: A TPB-Based SEM Approach

**DOI:** 10.1101/2025.09.01.673580

**Authors:** Xintong Chou, Hanyu Peng, Zhen Zhang, Huirong Ma, Zuobila Aimaitijiang, Xin Li, Liqun Wang, Hongyan Qiu

## Abstract

**Background:** Global fertility rates are declining, and despite China’s three-child policy, the number of women of childbearing age continues to shrink. Fertility intention, a key predictor of fertility behavior, is shaped by factors such as socioeconomic stress, family functioning, and psychological traits.

**Objective:** Grounded in the Theory of Planned Behavior, this study investigates how family functioning, marriage and fertility values, and fertility anxiety influence the fertility intentions of newlywed, childless young couples.

**Methods:** A cross-sectional survey of 1,292 newlywed, childless couples in Yinchuan, Ningxia (2024) assessed demographics, family functioning (FAD), marriage and fertility values (MFV), and fertility anxiety (FA). Multiple logistic regression and structural equation modeling (SEM) were used.

**Results:** Family functioning promoted intentions to have two or more children indirectly via positive communication and mediated by traditional MFV (44.4% of the total effect). Fertility anxiety showed no significant effect. Men preferred more children than women, and fertility intentions increased with age

**Conclusions:** Improving family communication and reinforcing traditional values may help boost fertility intentions. Findings offer insight into the family–psychological mechanisms shaping fertility decisions.

## Background

Global fertility has dropped from 4.98 in 1960 to 2.39 in 2020, and is expected to fall below the replacement level of 2.1 by 2035^1^. Low fertility has become a global challenge, disrupting population dynamics and worsening economic and social issues, especially in aging nations with shrinking workforces^2^. This trend of population decline has not improved, especially after the introduction of the three-child policy in China^3^. There were 9.54 million births in 2024, an increase of 0.52 million compared with 2023. However, the number of women of childbearing age (15-49), particularly those at the peak of their childbearing years (21-35), is still falling^4^.

In this perspective, fertility intentions have been the focus of research as an important indicator for understanding and predicting fertility behavior^5^. Fertility intentions refers to an individual’s expectation of having children, which is manifested in the time of willingness to have children, the ideal number of children, and the Ideal Gender of Children ^6^. Recent years have witnessed a noticeable decline in fertility intentions, driven by increasing socio-economic pressures and individuals’ growing emphasis on personal development. Although many people still uphold relatively high ideals regarding the number of children, studies have shown a gradual decrease in both the number of children they desire and actually intend to have ^3,7,8^. A growing body of research attributes this trend to multiple factors, including economic stress ^9^, women’s employment stress in the workplace^10^, family roles^11^, intergenerational family support^12^, gender ideology disparities^13^, fertility costs^14^, parenting stress^15^, etc. A Chinese meta-analysis further corroborates this perspective^16^.

In China, fertility patterns are often passed down across generations; more siblings in the family of origin are linked to stronger fertility intentions^17^. Research shows that parents significantly influence their children’s fertility plans. Family support helps shape fertility behavior.^18^, This influence is manifested through various forms of support, including care provision, financial assistance, and expressed preferences regarding the number of grandchildren^19^. The McMaster Model of Family Functioning (MMFF) views the family as a functional system encompassing problem-solving, communication, family roles, affective response, affective involvement, and behavior control. Failure to fulfill these functions may lead to clinical issues among family members^20^.While family-based fertility remains dominant, modern “love—economy— fertility” marriage models increase pressure on highly educated youth, leading them to delay childbearing and set higher expectations for children^21^. The Theory of Planned Behavior (TPB) posits that fertility intention (FI) derives not only from socio-economic rational calculations but is also shaped by individuals’ subjective emotional states and cognitive beliefs^22^, such as attitudes toward marriage and childbearing, including fertility values and fertility-related anxiety, which reflect underlying beliefs, values, and norms regarding marriage, family, and reproduction.

The Second Demographic Transition (SDT) framework provides a valuable lens through which to examine contemporary transformations in marriage and fertility patterns. Accompanying ongoing socio-economic development is a paradigmatic shift toward values centered on autonomy, self-actualization, and expressive individualism^23^.Traditional concepts such as lineage continuation are gradually losing their influence, particularly among younger, highly educated populations, leading to increasingly pluralistic attitudes toward fertility^24^. This shift in values—marked by the erosion of conventional family norms and the diversification of life trajectories—has contributed to notable demographic trends, including delayed marriage and childbearing, rising rates of childlessness, and the growing acceptance of alternative living arrangements. A study involving 1,492 Chinese university students found that only 9.68% expressed a clear desire to have children^25^, This shift in values—marked by the erosion of conventional family norms and the diversification of life trajectories— has contributed to notable demographic trends, including delayed marriage and childbearing, rising rates of childlessness, and the growing acceptance of alternative living arrangements. A study involving 1,492 Chinese university students found that only 9.68% expressed a clear desire to have children^26^.Nevertheless, human reproductive behavior is shaped by a complex interplay of factors. At present, research exploring the influence of evolving marriage, family, and fertility models on individual reproductive choices remains relatively limited. This study seeks to address this gap by constructing an integrated “individual–psychological–family” pathway model from micro-level psychological and familial perspectives, in order to better understand the mechanisms underlying reproductive decision-making in contemporary contexts.

By employing a structural equation model (SEM), this study aims to explore how family-related and psychological variables jointly influence young women’s fertility intentions. This integrated approach provides both theoretical and empirical insights into the multifaceted psychological mechanisms that underlie fertility decision-making among youth.

### Accordingly, the following research hypotheses are proposed

**H1:** Family functioning has a direct positive effect on the fertility intention of newly married, childless young couples.

(Family Assessment Device → Fertility Intention)

**H2:** Marriage and fertility values may mediate the relationship between family functioning and fertility intention.

(Family Assessment Device→ Marriage and Fertility Views → Fertility Intention)

**H3:** Fertility anxiety may mediate the relationship between family functioning and fertility intention.(Family Assessment Device→ Fertility Anxiety → Fertility Intention)

**H4:** Marriage and fertility values and fertility anxiety play a sequential mediating role in the relationship between family functioning and fertility intention.

(Family Assessment Device → Marriage and Fertility Views → Fertility Anxiety → Fertility Intention) See **Error! Reference source not found.**.

## Methods

### Study Design and Sample

This study adopts a simple random sampling method to select one of six marriage registration offices in Yinchuan, Ningxia, as the research site, with data collection conducted from March 15 to October 1, 2024. The study targets newlywed couples obtaining marriage certificates at the selected office. Inclusion criteria include young newlywed couples, while exclusion criteria involve cases in which either spouse has a history of remarriage or prior childbirth. The sample size was calculated using an epidemiological formula with a 0.05 margin of error, resulting in an estimated sample of 1,212 individuals (606 couples). To account for potential loss to follow-up, the final sample size was adjusted to 1,292 individuals (646 couples). Both quantitative and qualitative methods are employed, utilizing a self-designed questionnaire consisting of four sections: demographic characteristics, marriage and fertility views, intergenerational family relationships and support, and standardized measurement scales.

### Ethics and consent

This study was approved by the Medical Ethics Committee of Ningxia Medical University on December 12, 2023 (Approval No. 2023-057), and all participants (and/or their legal guardians) provided written informed consent in accordance with the Declaration of Helsinki and relevant guidelines and regulations

### Measurements and Instruments

The Theory of Planned Behavior, as the fundamental theoretical model for fertility intentions and behaviors, plays a significant role in predicting reproductive behaviors. According to this theory, fertility intentions are typically measured in terms of the timing, gender, and number of children. Moreover, individual background factors, including personal, social, and informational factors, influence these three direct determinants of fertility intentions. Based on the Theory of Planned Behavior and previous research, a survey questionnaire was developed to collect information on the following five key components.:

(1) **Demographic Characteristics**: Includes gender, age, residence (urban/rural), education (below/bachelor’s and above), employment, living with parents, and whether the individual is an only child. Household income is categorized into low (<5000 RMB), middle (5000-10,000 RMB), and high (>10,000 RMB) based on national statistics.
(2) **Fertility Anxiety Measurement**: **Subjective Anxiety**: Measured by the question, “Do you experience fertility anxiety?” The response options are: None, Slightly anxious, Somewhat anxious, Very anxious, and Uncertain. “None” and “Uncertain” were combined into the “no anxiety” category, while the other responses were categorized as “anxiety.”
(3) **Marry and Fertility Views:** Based on two questions: Views on love and marriage: 1) Love before marriage, 2) Can marry without love, 3) Only love, no marriage; Views on marriage and fertility: 1) Marriage’s main purpose is to have children, 2) Marriage and children are not linked, DINK acceptable, 3) Marriage is optional, can have children without marriage. Those selecting the first option for both questions are categorized as having continuous views, while others are classified as fragmented.
(4) **Measurement of Family Functioning**: This study uses the Revised Chinese Family Assessment Device (FAD), which includes five dimensions: Emotional Communication (reverse-scored), Positive Communication, Individualism (reverse-scored), Problem Solving, and Family Rules (reverse-scored), with 30 items in total. Emotional Communication evaluates the depth and style of emotional expression within the family. Positive Communication reflects active, open, and supportive interaction. Individualism measures whether family members prioritize self-interest over shared responsibilities. Problem Solving assesses how family issues are handled, while Family Rules looks at the presence of effective behavioral guidelines, such as household chores. A 4-point scale is used, from “not like my family” to “completely like my family,” with higher scores indicating better family functioning^20^, The Cronbach’s α coefficient in this study was 0.779.
(5) **Measurement of Fertility Intentions**: **Ideal number of children**: Measured by the question: “If there were no policies or restrictions, how many children would you ideally have?” Response options: “0, 1, 2 or more, doesn’t matter.” The “doesn’t matter” group is combined with “0” to represent undecided about having children. **Sex preference**: Measured by the question: “What is the ideal gender of your children?” Response options: “A boy, a girl, both, doesn’t matter.” **Age at first childbearing**: Measured by the question: “What is the ideal age for having your first child?” This is a continuous variable. Since ideal gender and age are influenced by the ideal number of children, this study incorporates the ideal number as a key dependent variable.

### Statistical analysis

Data were analyzed using SPSS 27.0 and MPLUS8.3. First, frequency and percentage were used to describe the characteristics of childless newlywed couples. Chi-square and ANOVA tests were applied to compare differences in ideal number of children (none, 1, 2 or more) across demographic variables. To identify factors influencing fertility intentions, multinomial logistic regression was conducted. Model 1 controlled for demographic variables and family functioning. Model 2 built upon Model 1 by adding fertility-related anxiety. Model 3 further included gender, age, and marriage and fertility values. All models used “no children” as the reference group, and odds ratios (OR) with 95% confidence intervals (CI) were calculated. Structural equation modeling (SEM) was performed using Mplus 8.3, and PowerPoint 2020 was used to visualize the model structure.

## Results

### The characteristics of study population

Table 1 shows the demographic characteristics of different ideal number of children groups. Age and gender significantly differed across groups. As the ideal number of children increased, average age also rose (p = 0.038). Men preferred having two or more children (p < 0.001). No significant differences were found in occupation, education, income, only-child status, or cohabitation with parents (p > 0.05). Higher proportions of individuals with a bachelor’s degree or higher, working in service industries, and earning under 5000 yuan were observed. Overall, ideal number of children differed mainly by age and gender.

**Table 1.** Sociodemographic Characteristics by Ideal Number of Children.

| Variable | Ideal Number of Children | | | $\chi^2/F$ | *p*-value |
| --- | --- | --- | --- | --- | --- |
|  | None | One | Two or More |  |  |
| Age (Mean ± SD) | 27.987±3.641 | 28.344±4.774 | 28.832±4.193 | 3.279 | 0.038* |
| Gender, n (%) |  |  |  |  |  |
| Male | 64(41.56) | 188(44.66) | 394(54.95) | 16.23 | <<br>0.001*** |
| Female | 90(58.44) | 233(55.34) | 323(45.05) |  |  |
| Occupation, n (%) |  |  |  |  |  |
| Farmer | 7(4.55) | 7(1.66) | 21(2.93) | 9.363 | 0.313 |
| Worker | 15(9.74) | 60(14.25) | 100(13.95) |  |  |
| Public sector employees<br>(government staff,<br>teachers, healthcare<br>professionals) | 37(24.03) | 107(25.42) | 155(21.62) |  |  |
| Business owners, private<br>enterprise employees | 21(13.64) | 71(16.86) | 123(17.15) |  |  |
| Service industry and<br>others | 74(48.05) | 176(41.81) | 318(44.35) |  |  |
| Educational Level, n (%) |  |  |  |  |  |
| Below Bachelor's degree | 20(12.99) | 80(19.00) | 126(17.57) | 2.834 | 0.242 |
| Bachelor's degree and<br>above | 134(87.01) | 341(81.00) | 591(82.43) |  |  |
| Monthly Income (RMB), n (%) |  |  |  |  |  |
| <5000 | 78(50.65) | 197(46.79) | 312(43.51) | 3.943 | 0.414 |
| 5000-10000 | 58(37.66) | 172(40.86) | 298(41.56) |  |  |
| >1000 | 18(11.69) | 52(12.35) | 107(14.92) |  |  |
| Only Child Status, n (%) |  |  |  |  |  |
| yes | 46(29.87) | 119(28.27) | 187(26.08) | 1.247 | 0.536 |
| no | 108(70.13) | 302(71.73) | 530(73.92) |  |  |
| Co-residence with Parents, n (%) |  |  |  |  |  |
| yes | 56(36.36) | 141(33.49) | 245(34.17) | 0.414 | 0.813 |
| No | 98(63.64) | 280(66.51) | 472(65.83) |  |  |

### The characteristics of Family Functioning, Fertility Views, Anxiety & Intentions

Table 2 summarizes the differences among groups with different ideal numbers of children. Participants with continuous marriage and fertility views were more likely to prefer having two or more children (*p* < 0.001). Fertility anxiety and its impact were most common in the one-child group (*p* < 0.001). Significant group differences were also found in family functioning, including emotional communication, positive communication, self-centeredness, problem solving, and family rules (*p* < 0.05), with the one-child group scoring highest in positive communication. Preferences for child gender varied significantly (*p* < 0.001), with the two-or-more group favoring “a boy and a girl,” and the no-child group favoring “no preference.” Expected childbearing age was earliest in the two-or-more group (*p* < 0.001).

**Table 2.** Group Differences by Ideal Number of Children.

| Variable | Ideal Number of Children | | | $\chi^2/F$ | *p*-value |
| --- | --- | --- | --- | --- | --- |
|  | None | One | Two or More |  |  |
| Marriage and Fertility View ,n (%) |  |  |  |  |  |
| Continuous | 12 (7.79%) | 57 (13.54%) | 206 (28.73%) | 55.535 | <0.001*** |
| Discontinuous | 142 (92.21%) | 364 (86.46%) | 511 (71.27%) |  |  |
| Fertility Anxiety, n (%) |  |  |  |  |  |
| No | 57 (37.01%) | 104 (24.70%) | 266 (37.10%) | 19.662 | <0.001*** |
| Yes | 97 (62.99%) | 317 (75.30%) | 451 (62.90%) |  |  |
| Impact of Fertility Anxiety, n (%) |  |  |  |  |  |
| No Impact | 45 (29.22%) | 104 (24.70%) | 258 (35.98%) | 16.063 | <0.001*** |
| Impact | 109 (70.78%) | 317 (75.30%) | 459 (64.02%) |  |  |
| Family Assessment Device ,(Mean ± SD) | | | | | |
| Total Score | 71.89 ± 11.32 | 72.62 ± 8.75 | 72.52 ± 10.31 | 0.315 | 0.730 |
| Emotional Communication | 20.03 ± 5.51 | 19.31 ± 5.53 | 20.18 ± 5.66 | 3.257 | 0.039** |
| Positive Communication | 11.12 ± 3.53 | 12.29 ± 3.52 | 11.12 ± 3.59 | 15.174 | <0.001*** |
| Self-Centeredness | 14.13 ± 4.40 | 14.24 ± 4.32 | 14.82 ± 4.51 | 3.094 | 0.046** |
| Problem Solving | 13.90 ± 3.91 | 14.60 ± 4.09 | 13.31 ± 4.17 | 13.215 | <0.001*** |
| Family Rules | 12.71 ± 3.24 | 12.18 ± 3.66 | 13.09 ± 3.62 | 8.561 | <0.001*** |
| sex preference |  |  |  |  |  |
| Have a bboy | 6(3.90) | 42(9.98) | 134(18.69) | 319.143 | <0.001*** |
| Have a girl | 14(9.09) | 131(31.12) | 115(16.04) |  |  |
| Boy and Girl | 27(17.53) | 66(15.68) | 356(49.65) |  |  |
| Donnot matter | 107(69.48) | 182(43.23) | 112(15.62) |  |  |
| Age at first<br>childbearing(Mean ± SD) | 28.36±2.76 | 28.50±3.16 | 27.51±3.05 | 15.843 | <0.001*** |

**Error! Reference source not found.** presents the basic distribution of fertility intentions. The left chart illustrates respondents’ preferences regarding the ideal number of children, with the majority (55.49%) indicating a preference for having two or more children, followed by 32.59% who preferred having only one child, and 11.92% expressing no intention to have children. The right chart displays preferences concerning the gender of children, where 34.75% of respondents reported no gender preference, 31.04% expressed a preference for having both a boy and a girl, 20.12% preferred having a girl, and 14.09% preferred having a boy.

**Error! Reference source not found.** shows the distribution of ideal childbearing age preferences among males and females, revealing distinct gender differences. Both genders have a primary peak at ages 29-30, with males at approximately 27% and females at 19%. This reflects societal recognition of this age range as ideal for childbearing. Females also show a secondary peak at ages 25-26, with a preference of around 15%, while males exhibit a slightly lower preference (12%) in the same age range, suggesting that women tend to plan for childbearing earlier. The male distribution is more spread out, with childbearing intentions ranging from 17-18 years (around –2%) to 41-42 years (around –3%), with relatively stable percentages in the 25-34 age group (–15% to –27%). In contrast, the female distribution is more concentrated, with around 70% of women showing a preference for childbearing between ages 25-36. After age 35, the preference sharply declines, and by age 40, it nearly vanishes, with a mere 0.5% at ages 49-50. Both males and females show a significant drop in childbearing intentions after age 45, with a preference of less than 0.5% at ages 49-50, indicating that societal norms view this as beyond the ideal childbearing window.

### Multinomial Logistic Regression of Ideal Number of Children (Fertility Intentions)

Table 3 presents the results of a multinomial logistic regression analysis examining the associations between the ideal number of children (fertility intentions) and key psychological and demographic factors. The outcome variable was categorized into three groups, with “no children” as the reference category; results are reported for the categories “having one child” and “having two or more children.”

**Table 3.** Multinomial Logistic Regression Analysis of Ideal Number of Children (Fertility Intentions) and Associated Factors.

| Variable | MODEL 1 |  |  | MODEL 2 |  |  | MODEL 3 |  |  |
| --- | --- | --- | --- | --- | --- | --- | --- | --- | --- |
|  | B | P | OR(95%CI) | B | P | OR(95%CI) | B | P | OR(95%CI) |
| **Having 1 Child** |  |  |  |  |  |  |  |  |  |
| Intercept | -1.428 | 0.037 |  | -1.422 | 0.039 |  | -0.406 | 0.662 |  |
| **FAD** |  |  |  |  |  |  |  |  |  |
| Emotional Communication | 0.012 | 0.538 | 1.012 (0.973,1.053) | 0.012 | 0.534 | 1.013(0.974,1.053) | 0.016 | 0.427 | 1.016(0.976,1.058) |
| Positive Communication | -0.03 | 0.322 | 0.970 (0.914,1.030) | -0.031 | 0.319 | 0.97(0.913,1.030) | -0.027 | 0.386 | 0.973(0.916,1.035) |
| Self-Centeredness | -0.039 | 0.132 | 0.962 (0.914,1.012) | -0.039 | 0.134 | 0.962(0.914,1.012) | -0.023 | 0.386 | 0.977(0.928,1.029) |
| Problem Solving | 0.04 | 0.149 | 1.041 (0.986,1.099) | 0.04 | 0.15 | 1.041(0.986,1.099) | 0.038 | 0.182 | 1.038(0.983,1.097) |
| Family Rules | 0 | 0.999 | 1 (0.938,1.066) | 0 | 0.996 | 1(0.938,1.066) | 0.007 | 0.842 | 1.007(0.943,1.075) |
| **Fertility Anxiety** |  |  |  |  |  |  |  |  |  |
| No |  |  |  | -0.009 | 0.963 | 0.991(0.688,1.428) | 0.31 | 0.121 | 1.363(0.922,2.106) |
| Yes |  |  |  | Reference |  |  | Reference |  |  |
| **Marriage and Fertility View** |  |  |  |  |  |  |  |  |  |
| Continuous |  |  |  |  |  |  | -1.546 | <0.001*** | 0.213(0.114,0.398) |
| Discontinuous |  |  |  |  |  |  | Reference |  |  |
| **Gender** |  |  |  |  |  |  |  |  |  |
| Boy |  |  |  |  |  |  | -0.402 | 0.037** | 0.669(0.458,0.977) |
| Girl |  |  |  |  |  |  | Reference |  |  |
| Age |  |  |  |  |  |  | -0.038 | 0.070 | 0.962(0.923,1.003) |
|  | B | P | OR(95%CI) | B | P | OR(95%CI) | B | P | OR(95%CI) |
| **Having 2 or more Child** |  |  |  |  |  |  |  |  |  |
| Intercept | -1.515 | 0.004 |  | -1.257 | 0.019 |  | -0.761 | 0.268 |  |
| FAD |  |  |  |  |  |  |  |  |  |
| Emotional Communication | 0.002 | 0.885 | 1.002 (0.975,1.030) | 0.005 | 0.736 | 1.005(0.977,1.033) | 0.007 | 0.611 | 1.007(0.979,1.036) |
| Positive Communication | 0.064 | 0.003** | 1.066 (1.022,1.111) | 0.058 | 0.006** | 1.060(1.017,1.105) | 0.06 | 0.005** | 1.062(1.018,1.108) |
| Self-Centeredness | 0.014 | 0.451 | 1.014 (0.979,1.050) | 0.009 | 0.619 | 1.009(0.974,1.045) | 0.018 | 0.318 | 1.019 (0.982,1.056) |
| Problem Solving | 0.037 | 0.061 | 1.037 (0.998,1.078) | 0.034 | 0.087 | 1.034(0.995,1.075) | 0.034 | 0.09 | 1.034 (0.995,1.076) |
| Family Rules | -0.04 | 0.08 | 0.960 (0.918,1.005) | -0.04 | 0.087 | 0.961(0.918,1.006) | -0.035 | 0.132 | 0.965 (0.922,1.011) |
| Fertility Anxiety |  |  |  |  |  |  |  |  |  |
| No |  |  |  | -0.495 | <0.001*** | 0.610(0.464,0.802) | -0.313 | 0.034** | 0.731 (0.547-977) |
| Yes |  |  |  | Reference |  |  | Reference |  |  |
| Marriage and Fertility View |  |  |  |  |  |  |  |  |  |
| Continuous |  |  |  |  |  |  | -0.808 | <0.001*** | 0.446 (0.318,0.624) |
| Discontinuous |  |  |  |  |  |  | Reference |  |  |
| Gender |  |  |  |  |  |  |  |  |  |
| Boy |  |  |  |  |  |  | -0.173 | 0.197 | 0.841(0.647,1.094) |
| Girl |  |  |  |  |  |  | Reference |  |  |
| Age |  |  |  |  |  |  | -0.02 | 0.187 | 0.980 (0.952,1.01) |
| R<sup>2</sup> | 0.041 |  |  | 0.053 |  |  | 0.104 |  |  |

For the intention to have one child, none of the five dimensions of the Family Assessment Device (FAD)—emotional communication, positive communication, self-centeredness, problem-solving, and family rules—showed statistically significant effects. However, marital and fertility views, measured as a continuous variable reflecting more non-traditional or delayed fertility attitudes, were significantly negatively associated with the intention to have one child (B = –1.546, p < 0.001, OR = 0.213, 95% CI (0.114, 0.398). This suggests that individuals with more non-traditional views are less likely to intend to have only one child. Gender also had a significant effect: males were more likely than females to intend to have one child (B = –0.402, p = 0.037, OR = 0.669, 95% CI (0.458, 0.977). Age was not statistically significant in this model (B = –0.038, p = 0.070, OR = 0.962, 95% CI (0.923, 1.003).

For the intention to have two or more children, positive communication emerged as a consistent and significant predictor. In Model 1, it had a significant positive effect (B = 0.064, p = 0.003, OR = 1.066, 95% CI (1.022, 1.111), which remained significant in Model 2 (B = 0.058, p = 0.006, OR = 1.060, 95% CI (1.017, 1.105) and Model 3 (B = 0.060, p = 0.005, OR = 1.062, 95% CI (1.018, 1.108). Fertility anxiety was negatively associated with the intention to have two or more children: individuals without fertility anxiety were more likely to express such intentions (Model 2: B = –0.495, p < 0.001, OR = 0.610, 95% CI (0.464, 0.802); Model 3: B = –0.313, p = 0.034, OR = 0.731, 95% CI (0.547, 0.977). Marital and fertility views again showed a significant negative association (B = –0.808, p < 0.001, OR = 0.446, 95% CI (0.318, 0.624), suggesting that individuals with less traditional views were less likely to prefer having two or more children. Neither gender (B = –0.173, p = 0.197, OR = 0.841, 95% CI (0.647, 1.094) nor age (B = –0.020, p = 0.187, OR = 0.980, 95% CI (0.952, 1.010) was a significant predictor in this category.

Across the three models, explanatory power (R²) increased with the inclusion of additional predictors: Model 1 (R² = 0.041), Model 2 (R² = 0.053), and Model 3 (R² = 0.104), with Model 3 providing the best fit.

### Structural Equation Modeling of Fertility Intentions

#### Correlation Analysis of Fertility Intentions

Table 4 shows that gender is positively correlated with marriage and fertility values (r = 0.172, p <0 .05) and fertility attitudes (r = 0.288, p <0 .05), and negatively correlated with family functioning (r = –0.120, p <0 .05) and fertility intentions (r = – 0.108, p <0 .05. Age is weakly positively correlated with fertility intentions (r = 0.071, p <0 .05). Family functioning is negatively correlated with marriage and fertility values (r = –0.156, p <0 .05) and fertility attitudes (r = –0.102, p <0 .05). Marriage and fertility values are significantly negatively correlated with fertility intentions (r = –0.202, p <0 .05). Fertility attitudes are positively correlated with marriage and fertility values (r = 0.193, p <0 .05), but show no significant correlation with fertility intentions.

**Table 4.** Descriptive Statistics and Correlation Matrix of Main Study Variables (N = 1292)

| Variable | M | SD | sex | age | FAD | MFV | FA | FI |
| --- | --- | --- | --- | --- | --- | --- | --- | --- |
| sex | 1.5 | 0.5 | 1 |  |  |  |  |  |
| age | 28.571 | 4.338 | -0.139** | 1 |  |  |  |  |
| FAD | 87.876 | 15.647 | -0.120** | 0.034 | 1 |  |  |  |
| MFV | 1.787 | 0.409 | 0.172** | -0.017 | -0.156** | 1 |  |  |
| FA | 0.67 | 0.471 | 0.288** | -0.04 | -0.102** | 0.193** | 1 |  |
| FI | 2.436 | 0.696 | -0.108** | 0.071* | 0.091** | -0.202** | -0.054 | 1 |

#### Analyzing the Serial Mediation Pathways of Fertility Intentions(SEM)

Regression analysis revealed that FAD significantly negatively predicted marital and fertility views (MFV), with B = –0.005, p < 0.001. MFV significantly positively predicted fertility attitudes (FA), with B = 0.177, p < 0.001. Additionally, FAD positively predicted FA, with B = 0.003, p = 0.032. When predicting the final outcome variable, fertility intention (FI), MFV showed a negative effect (B = –0.166, p = 0.012), and FA also negatively predicted FI (B = –0.721, p < 0.001). The direct effect of FAD on FI was not significant (B = 0.006, p = 0.064). Gender significantly predicted FI (B = –0.246, p < 0.001). See Table 5 for details.

**Table 5.**
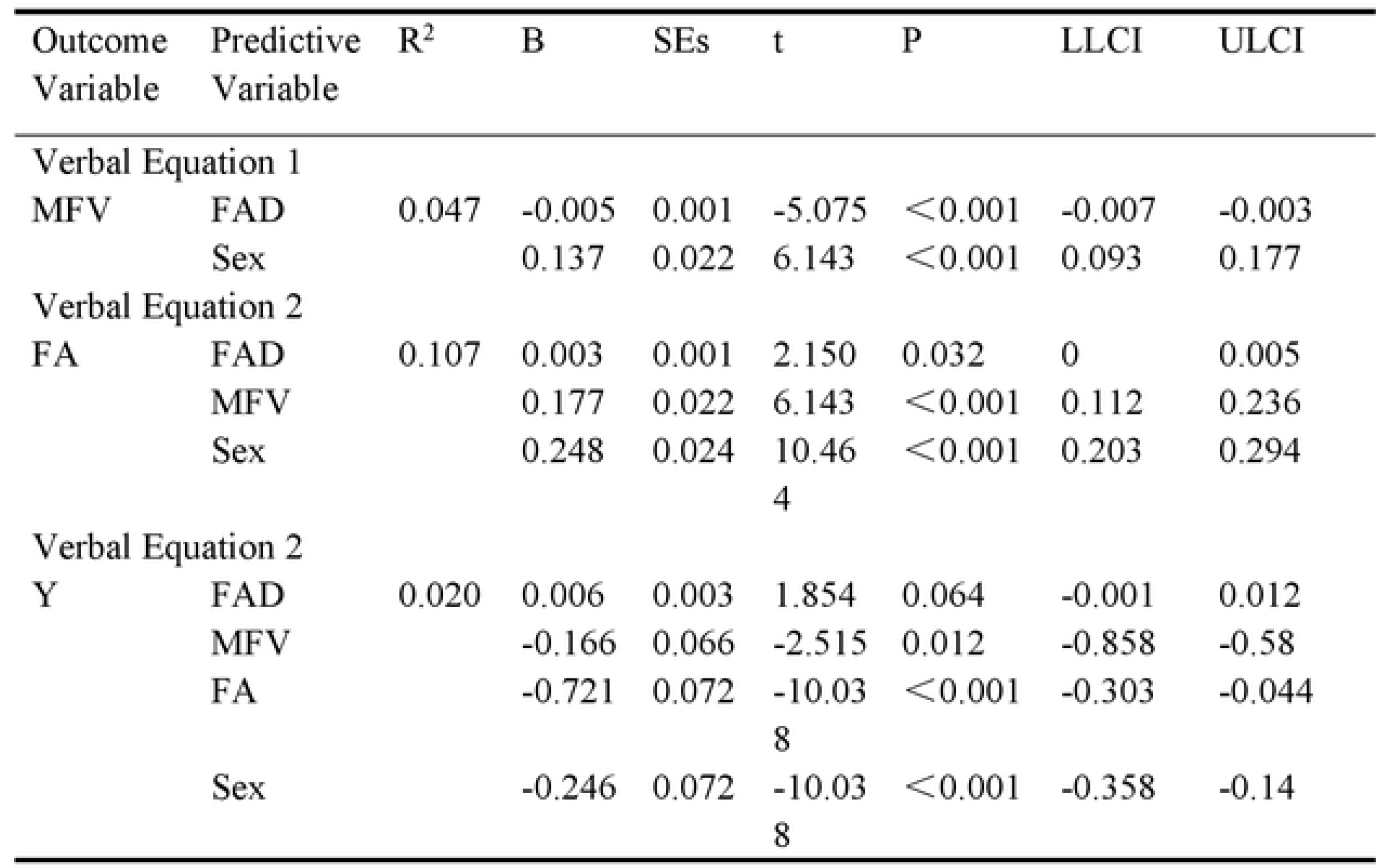
Serial Mediation Pathways with Bootslrap Estimates (N = 1,292) Using Maximum Likelihood.

As shown in Table 6, the total effect of FAD on fertility intention (FI) was significant (B = 0.009, SE = 0.003, 95% CI [0.004, 0.016]). The direct effect was not significant (B = 0.006, SE = 0.003, p = 0.064, 95% CI [–0.001, 0.012]). The total indirect effect was significant (B = 0.003, SE = 0.001, 95% CI [0.002, 0.005]), accounting for 33.3% of the total effect and 50.0% of the direct effect. Among the specific indirect pathways, only the FAD → MFV → FI path was statistically significant (B = 0.004, SE = 0.001, 95% CI [0.002, 0.006]), contributing 44.4% of the total indirect effect and 66.7% of the direct effect. The FAD → FA → FI and FAD → MFV → FA → FI paths were not significant (p = 0.120 and p = 0.055, respectively),( (see **Error! Reference source not found.** for details).

**Table 6.** Decomposition of Total, Direct, and Indirect Effects with Bootstrap Confidence Intervals (N = 1,292)

| Effect types | Effect | Boot SE | P | Boot LLCI | Boot ULCI | Ratio of indirect to total effect | Ratio of indirect to direct effect |
| --- | --- | --- | --- | --- | --- | --- | --- |
| Total effect | 0.009 | 0.003 | 0.004 | 0.003 | 0.016 |  |  |
| Direct effect | 0.006 | 0.003 | 0.064 | -0.001 | 0.012 |  |  |
| Total indirect effect | 0.003 | 0.001 | < 0.001 | 0.002 | 0.005 | 33.3% | 50.0% |
| FAD-MFV-FI | 0.004 | 0.001 | < 0.001 | 0.002 | 0.006 | 44.4% | 66.7% |
| FAD-FA-FI | 0.000 | 0.000 | 0.120 | -0.001 | 0.000 | 0% | 0% |
| FAD-MFV-FA-FI | 0.000 | 0.000 | 0.055 | 0.000 | 0.000 | 0% | 0% |

Table 7 reports the model fit indices for the serial mediation analysis controlling for gender, indicating that the model fits the data well. The chi-square (χ²) value is 0.000, with 0 degrees of freedom, indicating perfect fit between the model and the data; the RMSEA value is 0.000 (90% confidence interval = [0.000, 0.000]), showing perfect fit; the CFI and TLI values are both 1.000, indicating high fit between the model and the data; and the SRMR value is 0.000, further confirming the excellent fit. Additionally, the AIC is 6635.980, the BIC is 6713.439, the adjusted BIC is 6665.791, and the log-likelihood is –3302.990. These fit indices all support the model’s good fit to the data.

**Table 7.**
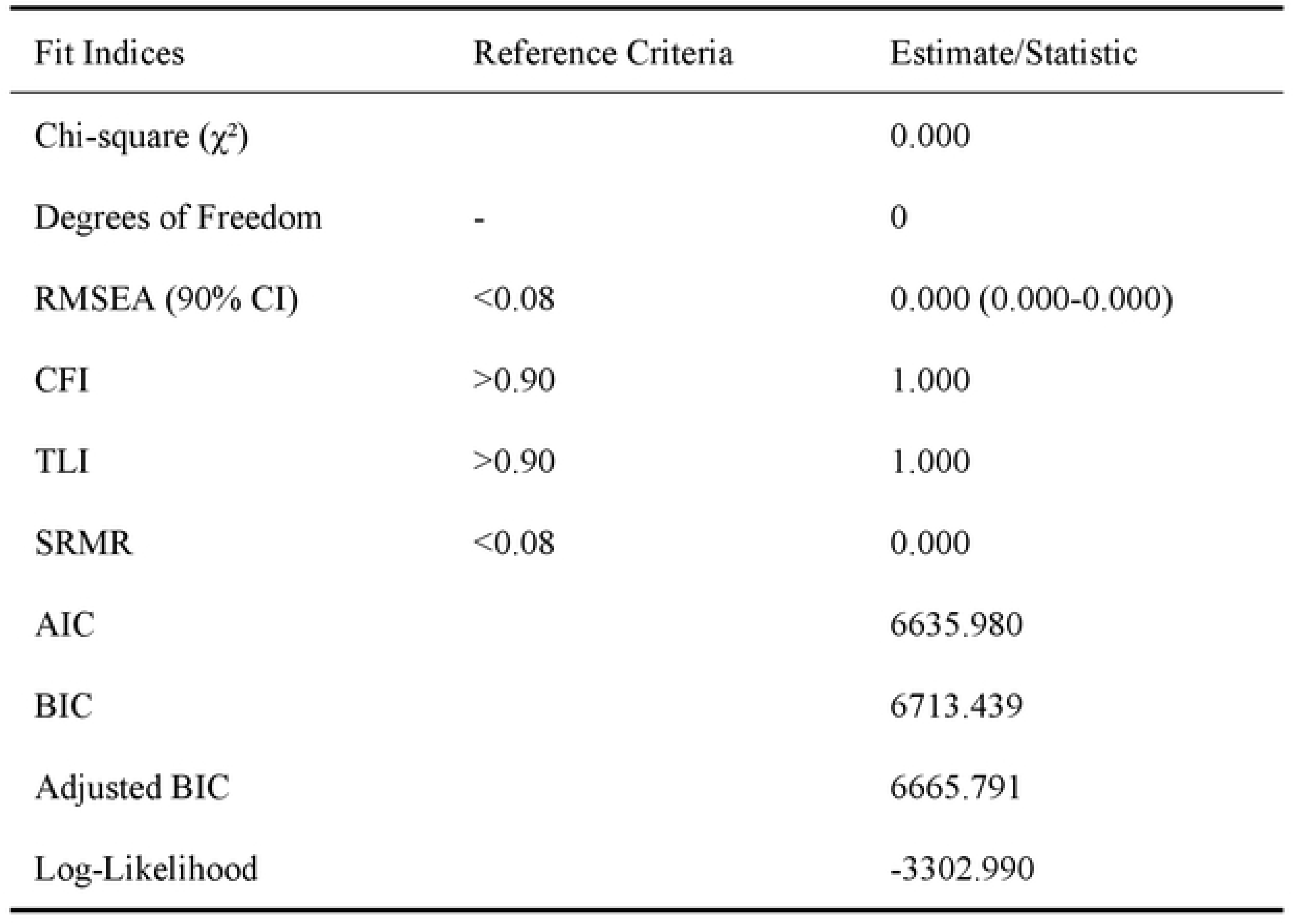
Model Fit Indices for the Serial Mediation Analysis Controlling for Gender (N = 1,292)

## Discussion

### The Role of FAD in Shaping Fertility Intentions

The results of this study are consistent with previous research, indicating that family functioning plays a critical role in shaping reproductive intentions and behaviors. In particular, effective family communication is a significant factor in promoting fertility intentions, especially when considering multiple children. Specifically, the “positive communication” dimension of family functioning was found to significantly predict the intention to have two or more children (B = 0.042, p < 0.01). Positive family communication has been shown to facilitate emotional support, reduce conflict, and enhance decision-making, all of which contribute to individuals’ confidence in their fertility decisions^27^.The study also identified a negative relationship between family functioning and marital and fertility views (MFV), consistent with the theoretical framework that suggests poor family functioning may lead to more negative or conservative fertility attitudes. Family functioning significantly predicted marital and fertility views in the negative direction (B = –0.005, p < 0.001). Prior research has indicated that individuals from dysfunctional family environments are more likely to internalize negative beliefs about marriage and childbearing, which may, in turn, influence their fertility intentions^28^. Furthermore, family functioning was positively associated with fertility anxiety (FA), suggesting that family dynamics can affect emotional preparedness for childbearing (B = 0.003, p = 0.032). Previous studies have indicated that individuals from high-conflict or emotionally unsupportive families often experience heightened anxiety about childbirth and childrearing, which may impede decision-making regarding family planning and contribute to a reluctance to have more children due to concerns about parenting^29–31^. It is noteworthy that the direct effects of family functioning on fertility intentions were not statistically significant in our models (B = –0.002, p = 0.112), which may suggest that other factors, such as socio-economic conditions, exert a more prominent influence on fertility decisions.

### Mediating Effects of Marriage-Fertility Views and Fertility Anxiety

In this study, we explored the mechanisms through which family functioning influences the fertility intentions of newlywed, childless individuals, using structural equation modeling. The results indicate that marital and fertility views (MFV) play a significant mediating role between family functioning (FAD) and fertility intentions (FI) (B = 0.004, SE = 0.001, 95% CI [0.002, 0.006]). This suggests that family functioning indirectly affects fertility intentions by influencing individuals’ marital and fertility views. Marital and fertility views, as individuals’ cognition and values regarding marriage and fertility, significantly shape their fertility decisions, further influencing the formation of fertility intentions. Therefore, marital and fertility views play an important role in fertility decisions, reflecting the profound impact of social and cultural backgrounds on individual fertility attitudes. Relevant studies have pointed out that factors such as social and cultural backgrounds^32^, educational levels^33^, and policy environments collectively affect individuals’ cognition and decisions regarding fertility^34^.However, the role of fertility anxiety in this study was relatively limited. Path analysis did not show significant results for the paths “FAD → fertility anxiety → fertility intention” or “FAD → marital and fertility views → fertility anxiety → fertility intention” (p = 0.120 and p = 0.055, respectively), indicating that fertility anxiety did not independently or jointly mediate the relationship between family functioning and fertility intention. This finding contrasts with previous studies, which have suggested that fertility anxiety may significantly influence fertility decisions, particularly in the context of multi-child policies and societal pressures^35^. For example, Jiang and Wu (2022) found a positive relationship between fertility anxiety and fertility decisions, especially in contexts where policies support multi-child families^36^. The increase in anxiety could lead individuals to hesitate about fertility. However, the results of this study may be due to the fact that in the specific cultural and social context of China, the influence of marital and fertility views is more pronounced. These views reflect more stable and long-term fertility attitudes, while fertility anxiety may have a more indirect or transient effect, reflecting short-term emotional fluctuations^37^. Thus, the impact of marital and fertility views on fertility intentions is more significant, while the effect of fertility anxiety on fertility intentions is relatively indirect or weak^38^. Future research could further explore the potential influence of fertility anxiety on fertility decisions in different social contexts and policy environments.

### Gender and Age Differences in Fertility Preferences

This study found no significant differences in occupation, education, income, whether the individual is an only child, or co-residence with parents (p > 0.05), which is consistent with previous research^3^. Women exhibit a more balanced distribution in their choice of the number of children, meaning that women show less variation in their fertility intentions across different child number categories, while men tend to prefer having a larger number of children. This gender difference suggests the potential influence of social, cultural, and gender roles on fertility decision-making. In many cultures, especially those with strong traditional values, men are typically seen as the economic providers for the family, and traditionally they have been expected to maintain and expand family size. As a result, men’s fertility intentions may lean more toward supporting the idea of larger families, although this trend may be gradually changing in modern society^37^. However, this difference may also reflect a difference in the way men and women perceive their responsibilities for child-rearing, with men perhaps relying more on their spouses or female family members to bear the brunt of child-rearing responsibilities, thus leading them to have higher expectations regarding the number of children^39^. On the other hand, as age increases, the ideal number of children rises significantly (p = 0.038). This trend suggests that older individuals are more inclined to support larger families. For example, the average age of individuals in the “no children” group was 27.99 years, in the “one child” group was 28.34 years, and in the “two or more children” group was 28.83 years. This phenomenon may reflect the fact that as individuals age, they become more aware of the timing of reproduction and the planning of future family life. Older individuals may face the pressure of biological timing and may feel a stronger need to achieve their fertility goals within a limited period^17^. Therefore, they are more likely to plan for having more children to ensure family stability and the continuation of their lineage. Additionally, older individuals often have a more solid economic foundation and a more stable career, which may lead them to feel that raising multiple children is more feasible under stable living conditions^40^.

## Conclusion

This study examines how family functioning influences the three-child fertility intentions of newlywed, childless young couples. H1 was not supported, showing that family functioning does not directly affect fertility intentions but works through mediating pathways. H2 was partially supported, with marriage and fertility values fully mediating the relationship, emphasizing the role of traditional family values in intergenerational transmission. H3*was not supported, as fertility anxiety did not significantly mediate the relationship, indicating that fertility decisions in a collectivist culture are more influenced by stable value systems than short-term emotional fluctuations. H4was not supported, as no sequential mediating effect of marriage values and fertility anxiety was found. In conclusion, family functioning impacts fertility intentions through marriage and fertility values.

### Implications for Policy and Family-Based Interventions

The findings of this study offer practical implications for optimizing fertility policies and designing targeted interventions. Family functioning plays a key role in shaping individuals’ marital and fertility views and intentions, suggesting that improving family dynamics—especially positive communication—should be a priority in fertility-related programs. Supportive family interactions can enhance individuals’ confidence and motivation in making reproductive decisions. Therefore, policy efforts should include premarital education, communication skills training, and psychological support to strengthen emotional bonds within families and foster positive fertility attitudes. The mediating role of marital and fertility views also indicates that interventions should go beyond financial incentives or direct policy measures. Instead, shaping values and perceptions about marriage and fertility is essential. Public education through diverse media platforms can help correct negative beliefs—such as concerns about parenting stress or high child-rearing costs—and reinforce the social value of family, thereby strengthening fertility intentions at the cognitive level.

### Limitations and Future Research Directions

The sampling scope of this study is limited to a single marriage registration institution in a city in Northwest China, which poses limitations on the geographical and cultural representativeness of the sample. It may not fully reflect the impact of urban-rural differences and regional cultural characteristics on fertility intentions across the country. The study design employs a cross-sectional survey, which cannot capture the dynamic evolution of fertility intentions among newlywed couples. Additionally, the exclusion of remarried couples and those with children may introduce bias when generalizing the conclusions to a broader reproductive-age population.

## Acknowledgements

We acknowledge all respondents who showed great patience in answering the questionnaires. We sincerely thank all those who have helped us.

## Funding info

This work was supported by the General Project of the National Social Science Foundation of China (Grant No. 23BRK036).

## Data available statement

Due to the continuity of the research project, the datasets generated and analyzed during the current research period are not made public, but can be obtained from the corresponding authors upon reasonable request.

---

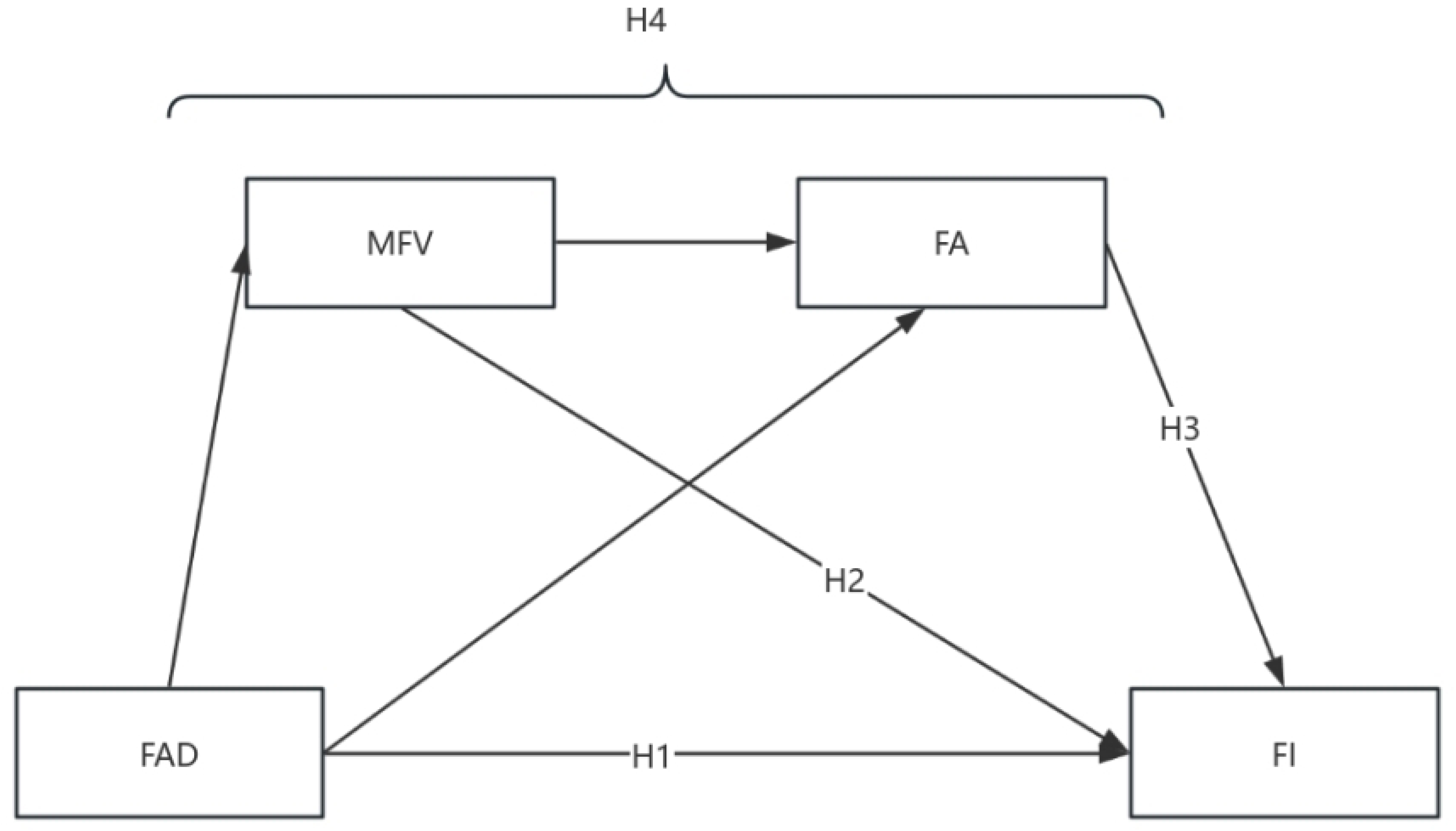
Figure 1.

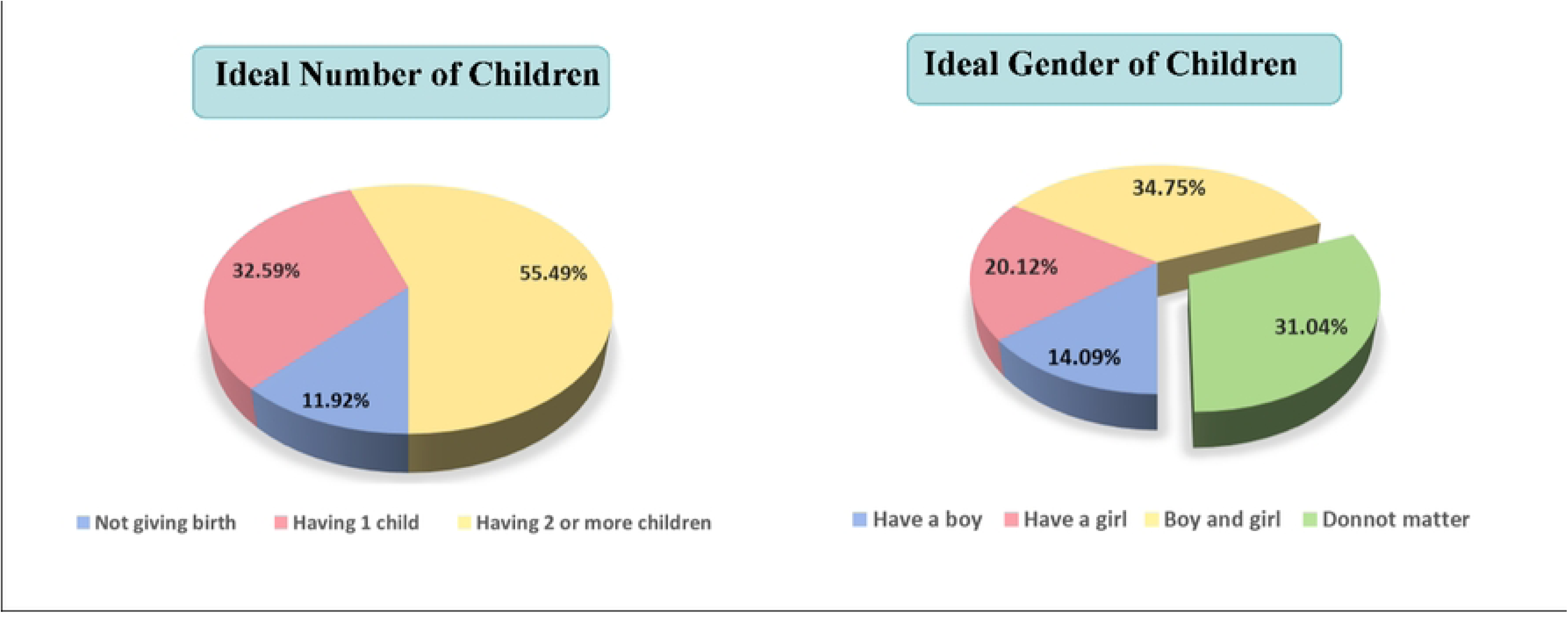
Figure 2.

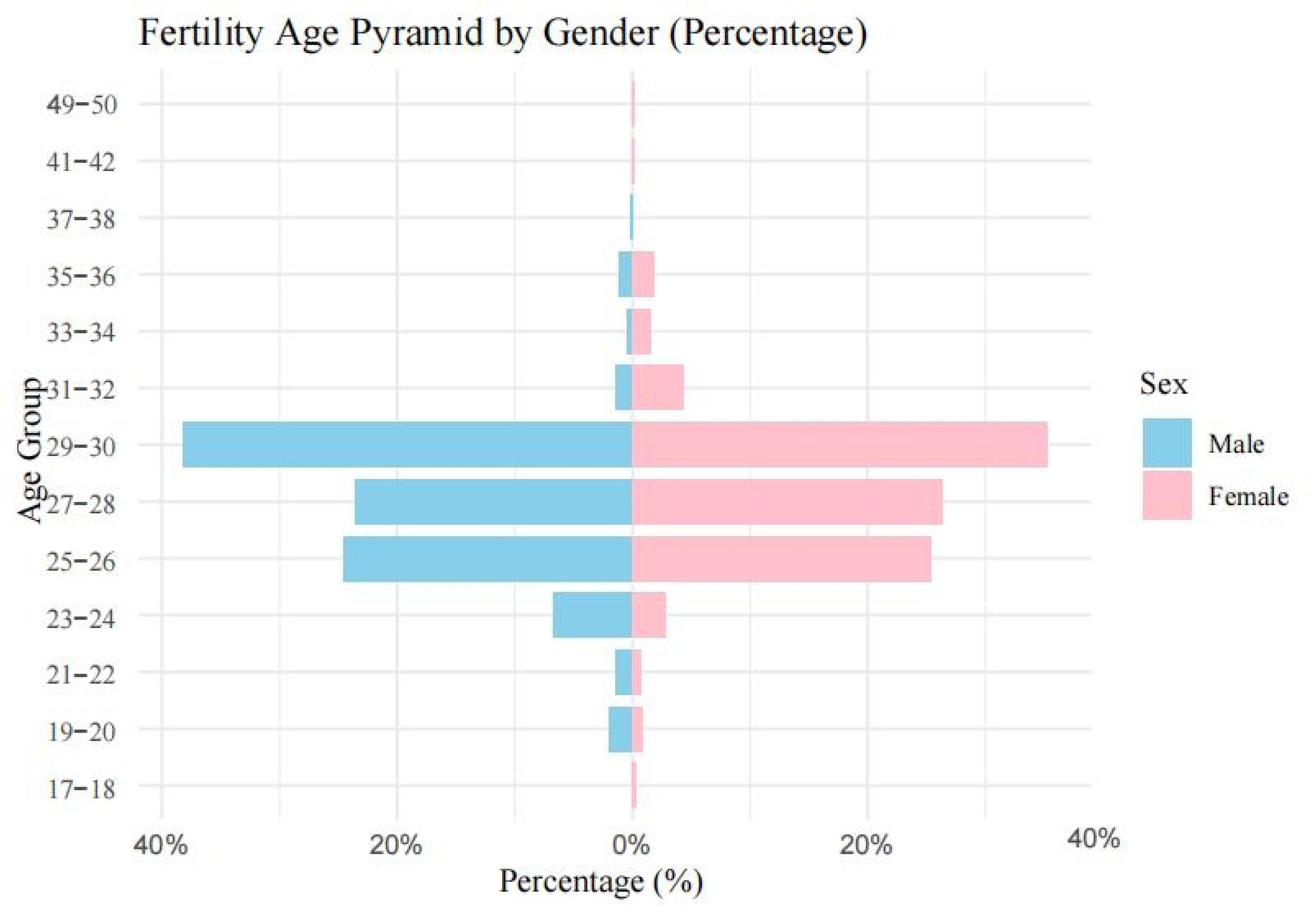
Figure 3.

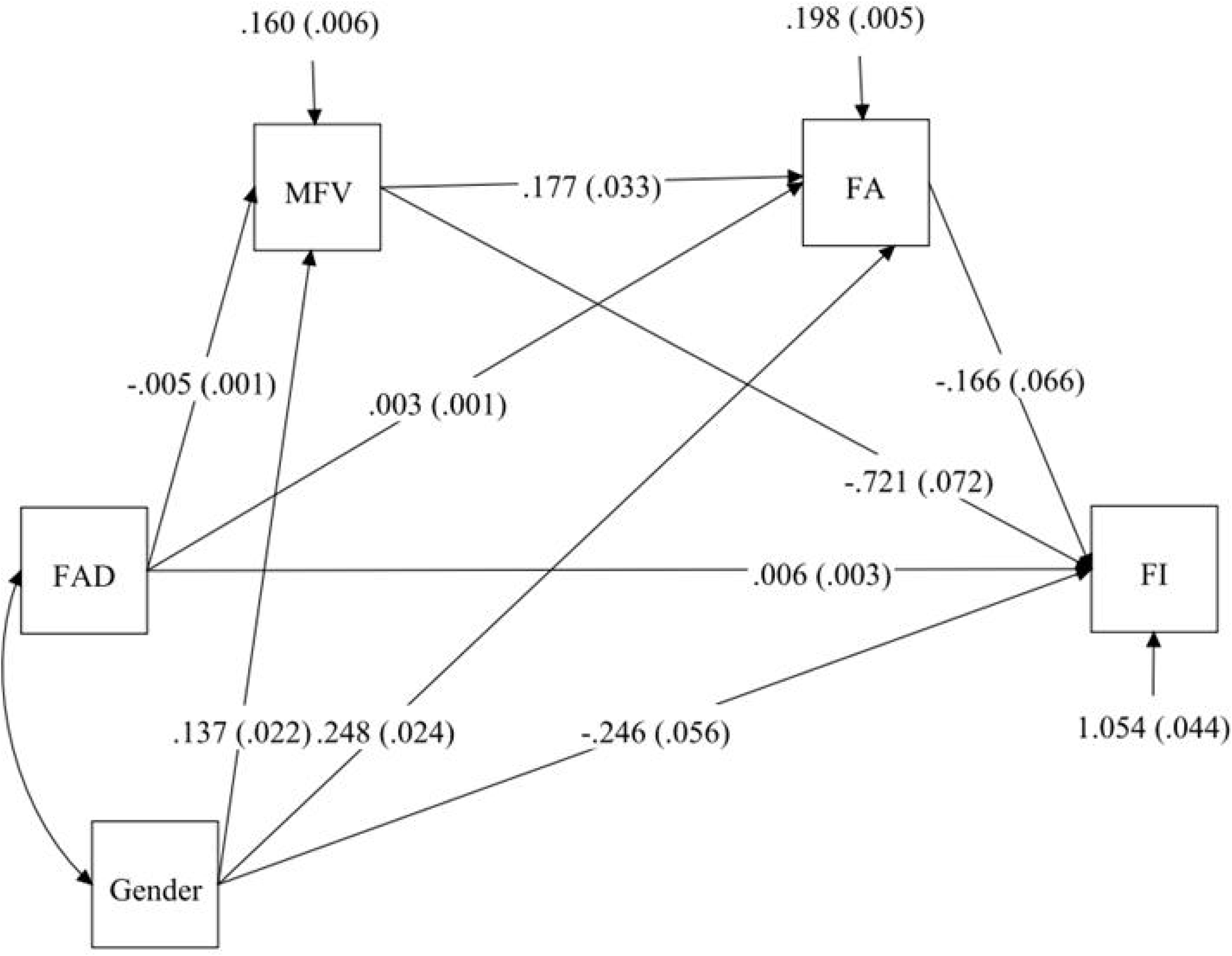
Figure 4.

